# Exomer cooperates with Ryh1^RAB6^ in regulating TORC2 activity

**DOI:** 10.1101/2025.07.15.664877

**Authors:** Esteban Moscoso-Romero, Sandra Moro, Kazuhiro Shiozaki, M.-Henar Valdivieso

**Affiliations:** Departamento de Microbiología y Genética, Universidad de Salamanca, Salamanca, Spain; Instituto de Biología Funcional y Genómica (IBFG), Consejo Superior de Investigaciones Científicas (CSIC), Salamanca, Spain; Nara Institute of Science and Technology, Ikoma, Japan; Department of Microbiology and Molecular Genetics, University of California, Davis, CA, USA

**Keywords:** Exomer, Ryh1, RAB6, *Schizosaccharomyces pombe*, stress, TORC2, yeast

## Abstract

Exomer is a trans-Golgi protein complex involved in multiple biological processes, including lipid homeostasis and stress response. We have found that in fission yeast, the absence of exomer leads to a defect in the recovery of “target of rapamycin complex 2” (TORC2) activity in response to high concentrations of KCl, but not to sorbitol, which indicates that the mutants are impaired in their ability to respond to salt stress. Changes in lipid homeostasis did not suppress this defect. Ryh1^RAB6^ is a Rab GTPase that resides in the Golgi and endosomes and facilitates or stabilizes the interaction between TORC2 and its substrate, the AGC kinase Gad8. We have found genetic and functional interactions between exomer and Ryh1^RAB6^. The recovery of TORC2 activity in response to KCl requires Ryh1^RAB6^ and exomer, and the localization of Ryh1^RAB6^ is aberrant in exomer mutants. Co-immunoprecipitation experiments indicate that the interaction between TORC2 and Gad8 is weaker in the absence of exomer. We propose that exomer regulates TORC2 by facilitating appropriate Ryh1 localization.

## INTRODUCTION

Exomer is a protein complex described as essential for delivering transmembrane proteins from the Golgi to the plasma membrane (PM) because, in *Saccharomyces cerevisiae* exomer mutants, these proteins stall in the Golgi (Santos et al., 1997; Santos and Snyder, 2000; Sanchatjate and Schekman, 2006; Trautwein et al., 2006; Wang et al., 2006; Barfield et al., 2009; Ritz et al., 2014). Nevertheless, exomer mutants exhibit some defects that cannot be ascribed to the retention of PM proteins in the Golgi (Sanchatjate and Schekman, 2006; Trautwein et al., 2006; Anton et al., 2017). The *Schizosaccharomyces pombe* exomer consists of Cfr1 and Bch1, which depend on each other for their localization in the Golgi (Hoya et al., 2017; Anton et al., 2018). In this organism, no proteins are known to stall in the Golgi in exomer mutants; in *S. pombe*, the absence of exomer leads to mild defects in multiple processes, including cell fusion during mating, cytokinesis, endomembrane system integrity, and protein trafficking (Cartagena-Lirola et al., 2006; Hoya et al., 2017). Additionally, these mutants are sensitive to high concentrations of KCl. Alongside this sensitivity, there is an aberrant calcium and potassium homeostasis, as well as an abnormal distribution of potassium and calcium channels and pumps. (Hoya et al., 2017; Moro et al., 2021). Finally, the exomer mutants exhibit defects in lipid homeostasis. In particular, they have a defect in the transport of phosphatidylinositol 4-phosphate (PI4P) from the Golgi to the PM, resulting in reduced association of Pck2 with the membrane and an inefficient Cell Integrity Pathway (CIP) response to KCl and sorbitol (Moscoso-Romero et al., 2024). The mutants are defective in the CIP response to both KCl and sorbitol; nevertheless, they are sensitive to KCl and grow efficiently on 1.2 M sorbitol, suggesting that the absence of exomer may lead to defects in additional signaling pathways that contribute to KCl sensitivity.

Target of Rapamycin (TOR) kinases are members of the phosphoinositide 3-kinase family that play a role in stress signaling and cellular homeostasis. TOR kinases associate with other proteins to form two distinct structural and functional complexes, TORC1 and TORC2 (reviewed in Eltschinger and Loewith, 2016; Morozumi and Shiozaki, 2021; Emmerstorfer-Augustin and Thorner, 2023). TORC1 is the best-characterized of the TOR complexes. It binds to the cytosolic leaflet of the lysosome/vacuole membrane and senses the cellular nutritional status. TORC1 activation promotes the uptake of nutrients and the synthesis of proteins, lipids, and nucleotides, while TORC1 inactivation promotes catabolism and autophagy. On the other hand, TORC2 is primarily located in the cytosolic leaflet of the PM. It responds to changes in PM tension and plays a role in regulating the actin cytoskeleton and PM-associated processes, such as endocytosis and PM integrity (Eltschinger and Loewith, 2016; Morozumi and Shiozaki, 2021; Emmerstorfer-Augustin and Thorner, 2023).

Yeast genomes encode two related TOR kinases, Tor1 and Tor2. In *S. pombe*, Tor2 is the catalytic subunit of TORC1, and Tor1 is that of TORC2. Tor1 associates with Ste20 (a homolog of the human protein RICTOR and the *S. cerevisiae* protein Avo3), Sin1 (*S. cerevisiae* Avo1), Bit61 (structurally related to human PROTOR/PRR5), and Lst8/Wat1, which also participates in TORC1 (Hayashi et al., 2007). The primary substrate of *S. pombe* TORC2 is Gad8, which is an AGC kinase homologous to human AKT1/AKT2 and *S. cerevisiae* YPK1/YPK2 (Matsuo et al., 2003; Ikeda et al., 2008). Gad8 undergoes multiple regulatory phosphorylation events, one of which is carried out by TORC2 on the serine at position 546 (henceforth referred to as Gad8^S546^) and can serve as a readout for TORC2 activity (Matsuo et al., 2003; Tatebe et al., 2010; Halova et al., 2013; Du et al., 2016). The components of the TORC2-Gad8 signaling pathway are found in both the nucleus and on the cell surface (Tatebe et al., 2010; Cohen et al., 2016). This pathway participates in sexual development as well as the response to different sources of stress, including DNA damage or replication stress, glucose starvation, high or low temperatures, oxidative stress, and osmotic stress (Weisman et al., 1997; Kawai et al., 2001; Weisman and Choder, 2001; Weisman et al., 2007; Schonbrun et al., 2013; Cohen et al., 2014; Weisman et al., 2014; Cohen et al., 2016).

*S. pombe* Ryh1, a homolog of human RAB6 (Hengst et al., 1990), is an upstream regulator of TORC2. Ryh1 deletion and inactivation result in phenotypes similar to those of deletion of the TORC2 components, including a drastic reduction of Gad8^S546^ phosphorylation (He et al., 2006; Tatebe et al., 2010). Ryh1^RAB6^ absence does not cause changes in TORC2 and Gad8 localization, so it has been proposed that Ryh1^RAB6^ facilitates the physical interaction between Tor1 and Gad8 (Tatebe et al., 2010). Ryh1^RAB6^ localizes in the Golgi/endosomes and participates in vesicle trafficking and the maintenance of vacuolar integrity independently of its function in TORC2 regulation (He et al., 2006).

The fact that both exomer and TORC2 mutants are sensitive to KCl (Kawai et al., 2001; Weisman and Choder, 2001; Matsuo et al., 2003; Ikeda et al., 2008; Tatebe et al., 2010; Moscoso-Romero et al., 2024) prompted us to investigate the existence of a functional relationship between exomer and TORC2.

## MATERIALS AND METHODS

### Strains and growth conditions

The yeast strains used in this work are listed in Table 1. The *cfr1Δ* and *bch1Δ* mutants were used interchangeably as exomer mutants because in *cfr1Δ,* Bch1 does not associate with the Golgi, and vice versa (Hoya et al., 2017). As a result, none of the mutants bear a functional exomer. Although each protein might participate in some exomer-independent function(s) by itself, we have never observed a difference in their phenotypes (Hoya et al., 2017; Moscoso-Romero et al., 2024, and results not shown). In our early work, we used the *cfr1::his3+* mutant, which we combined with other mutants or strains bearing tagged protein by genetic crosses, because homologous recombination at the *cfr1+* locus is inefficient. Later, with the use of the PCR-based technology to create antibiotic-resistance cassettes (Bähler et al., 1998), we found that *bch1::KAN*, *bch1::NAT*, and *bch1::HPH* mutants were efficiently produced, and that they exhibited identical phenotypes to *cfr1::his3+*. YES medium (0.5% yeast extract, 3% glucose, 225 mg /ml adenine, histidine, leucine, uracil and lysine hydrochloride, 2% agar, pH 5.6) was used for the growth and maintenance of *S. pombe* strains, as described (Moreno et al., 1991; Forsburg and Rhind, 2006). When required, geneticin (G418, Formedium), hygromycin B (Formedium), and nourseothricin (Werner BioAgents) were used at 120, 200, and 50 µg/ml, respectively. When necessary, miconazole (Sigma, Miconazole nitrate salt, n° M3512-1G) was added to the cultures at 6 µg/ml, final concentration, 1 hour before adding KCl. Samples for biochemical and microscopy analyses were taken from cultures growing exponentially in YES liquid medium at 28°C. To induce stress, samples were collected at various times after adding 1 M KCl (Merck, n° 1049361000) to the cultures. To analyze the response to hyperosmotic stress, cells were collected by filtration, and the filters were introduced into flasks with YES medium supplemented with 1.2 M sorbitol (Sigma, n° S1876), and incubated. The control sample (0’) was collected immediately before KCl was added, and before cells were transferred to sorbitol-supplemented YES. Drop-test analyses were performed as described (Hoya et al., 2017).

### Microscopy

For all the experiments involving confocal live-cell imaging, a NIKON Ti2-E confocal “spinning disk” microscope (100×/1.45 Oil Plan Apo objective), equipped with an Andor Dragonfly system and a sCMOS Sona 4.2B-11 camera, was used. Images were processed using deconvolution in the Fusion software from Andor. Stacks of three Z-series sections corresponding to the cell middle were acquired at 0.2-μm intervals. The images are average (AVG) projections. mCherry-FYVE(EEA1), a probe specific for PI3P, was used as a marker for the prevacuolar compartment (PVC). Krp1 (kexin, type I membrane endopeptidase) fused to the fluorescent protein tdTomato was used as a marker for the Golgi. Typically, samples were collected by centrifugation (1 minute at 3000 rpm), spotted on a slide, and photographed. To determine the fluorescence intensity of the proteins, the Fiji particle analysis tool (ImageJ software, National Institutes of Health) was used. We used the particle Fiji analysis tool to estimate the number of fluorescent dots per cell; a size threshold of 0.2 µm^2^ was established to eliminate background, and the points were counted manually. To determine the presence of Ste20^RICTOR^-GFP on the cell surface, fluorescent dots were scored from the poles and sides of cells that were not undergoing cytokinesis. To estimate the fluorescence intensity of intracellular structures, the Fiji particle analysis tool was used after defining a threshold. For co-localization analyses, stacks of three 0.20-µm Z-sections of the cell middle were acquired, and the central planes of the stacks from both channels were merged.

### Protein methods

For western blots, proteins in cell extracts were precipitated with trichloroacetic acid (TCA) and subjected to western blot analyses, as described previously (Moscoso-Romero et al., 2024). Cells growing exponentially in 30 ml of YES were collected by centrifugation (900 x g), washed with 1 ml of cold 20% TCA, and resuspended in 50 µl of the same solution. 500 µl of glass beads (Braun Biotech International) were added, and the cells were broken in a cold Fast Prep FP120 (Savant Bio101) using three 16-second pulses (speed 6), with 5-minute incubations on ice between pulses. 400 µl of cold 5% TCA was added to the tube, which was vortexed to wash the beads. Cell extracts were transferred to a clean tube and centrifuged for 10 minutes at 900 x g, at 4°C. The pellets were resuspended in 2% sodium dodecyl sulfate (SDS)/0.3 M Tris Base. The protein concentration was determined using the Bradford protein assay reagent (Bio-Rad, n° 500-0006). Equal amounts of protein were boiled in the presence of Laemmli sample buffer (50 mM Tris-HCl [pH 6.8], 1% SDS, 143 mM β-mercaptoethanol, 10% glycerol) for 5 minutes. The samples were subjected to polyacrylamide gel electrophoresis (PAGE 8% gels, 40% acrylamide/Bis solution 29:1. Bio-Rad, n° 161-0146), transferred to polyvinylidene difluoride (PVDF) membranes (Immobilon-P de PVDF. 0,45 micras. Millipore, n° IPVH00010), and incubated with non-fat dried milk to block nonspecific protein binding (Nestlé; 5% in TBST: 0.25% Tris-HCl, 0.9% NaCl, 0.25% Tween 20, pH 6.8). To assess the level of Gad8 phosphorylation in S546, the primary antibodies anti-Gad8^S546^ and anti-Gad8 (Tatebe et al., 2010; 1:4000) were used. The secondary antibody was horseradish peroxidase (HRP)-conjugated anti-rabbit IgG (Sigma, clone RG-96, 1:10000). To assess the level of Ste20^RICTOR^-GFP, the primary antibody was anti-GFP (JL8, BD Living Colors, 1:3000), and the secondary antibody was horseradish peroxidase (HRP)-conjugated anti-mouse IgG (Bio-Rad, n° 170-6515, 1:10000). The molecular weight standard was PageRuler Plus Prestained Protein Ladder (Thermo Scientific, n° 26620). Chemiluminescent signal was generated using the Western Bright ECL detection kit (Advansta, n° K-12045-D20) and detected on a Vilber Fusion FX system (Vilber GmbH), which captures chemiluminescence until the best-fitting signal is recorded.

Co-immunoprecipitation was performed as described (Yanguas and Valdivieso, 2021). Cells were washed with STOP Buffer (10 mM ethylenediaminetetraacetic acid [EDTA], 154 mM NaCl, 10 mM NaF, and 10 mM NaN_3_) and with Lysis Buffer (20 mM 4-(2-hydroxyethyl)-1-piperazineethanesulfonic acid [HEPES], pH 7.5; 150 mM monosodium glutamate, 0,25% Tween-20, 1µl/ml aprotinin, 1µl/ml leupeptin/pepstatin, 10µl/ml PMSF and phosphatase inhibitors; Cell signaling, n° 5872), and broken in a cold Fast Prep FP120 (Savant Bio101) in the same buffer using two 16-second pulses (speed 5), with 5-minute incubations on ice between pulses. Cell debris was removed by 30 minutes of centrifugation at 17000 x g. The protein concentration was determined using the Bradford protein assay reagent (Bio-Rad). 90 μg of the cleared cell lysates were boiled in the presence of Laemmli sample buffer and used as “whole cell extract”. 3 mg of lysates were combined with 50 μl of anti-GFP μMAC magnetic beads (Miltenyi Biotec, n° 130-091-125) and incubated in a tube rotator for 30 minutes at 4°C. The lysates were then applied in columns (Miltenyi Biotec, n° 130-042-701) previously prepared with 200 µl of Lysis Buffer, following the manufacturer’s instructions, washed 4 times with 200 µl of Wash Buffer 1 (150 mM NaCl, 1% Ecosurf EH-9, 0.5 M sodium deoxycholate, 0.1% SDS, 50 mM Tris-HCl pH 8.0), and washed one time with Wash Buffer 2 (20 mM Tris-HCl pH 7.5). To elute the proteins, 20 µl of 95°C hot Elution Buffer (50 mM Tris-HCl pH 6.8, 50 mM DTT, 1% SDS, 1 mM EDTA, 0.005% bromophenol blue, 10% glycerol) was applied to the column; after a 5-minute incubation time, 50 additional µl of hot Elution Buffer was added. Samples were boiled in Laemmli sample buffer and loaded into 8% polyacrylamide gels as described above. For quantifications, Fiji software was used to estimate the intensity of each band. Then, the value for the Gad8-HA coimmunoprecipitated was related to the HA total extracts in the same sample.

### Statistical analyses

Data quantification was performed on the results of a minimum of three experiments. To calculate the level of TORC2 activity in a sample, the intensity of phosphorylated Gad8 (Gad8^S546^) and total Gad8 bands in the western blot was estimated using Fiji. The ratio between the Gad8^S546^ and Gad8 band intensity was calculated. The mean, standard deviation, and statistical significance are depicted in bar graphs below the blot images in each Figure. For microscopy data, a minimum of 120 cells/particles from three different images were scored in each experiment. All the data were initially analysed with Excel, and then visualized and statistically analyzed with GraphPad Prism. In each case, the correction test suggested by GraphPad Prism was used to determine the statistical significance of the differences.

## RESULTS

### Exomer mutants are defective in the recovery of TORC2 activity after salt shock

To investigate the involvement of exomer in TORC2 signaling, we determined Gad8^S546^ phosphorylation in the wild-type control (WT in Figure 1A) and the exomer mutants *cfr1Δ* and *bch1Δ* exposed to 1 M KCl. We found that the level of activity in all strains was lower 15 minutes after the salt was added to the medium than in untreated cultures (0 minutes). After 90 minutes of incubation, activity was strong in the WT but weak in the mutants. The defect in reactivating the pathway was observed even after 3 hours of incubation (Figure 1B), indicating that the defect was not transient. Thus, exomer mutants are defective in the reactivation of TORC2 after high KCl stress.

**Figure 1.**
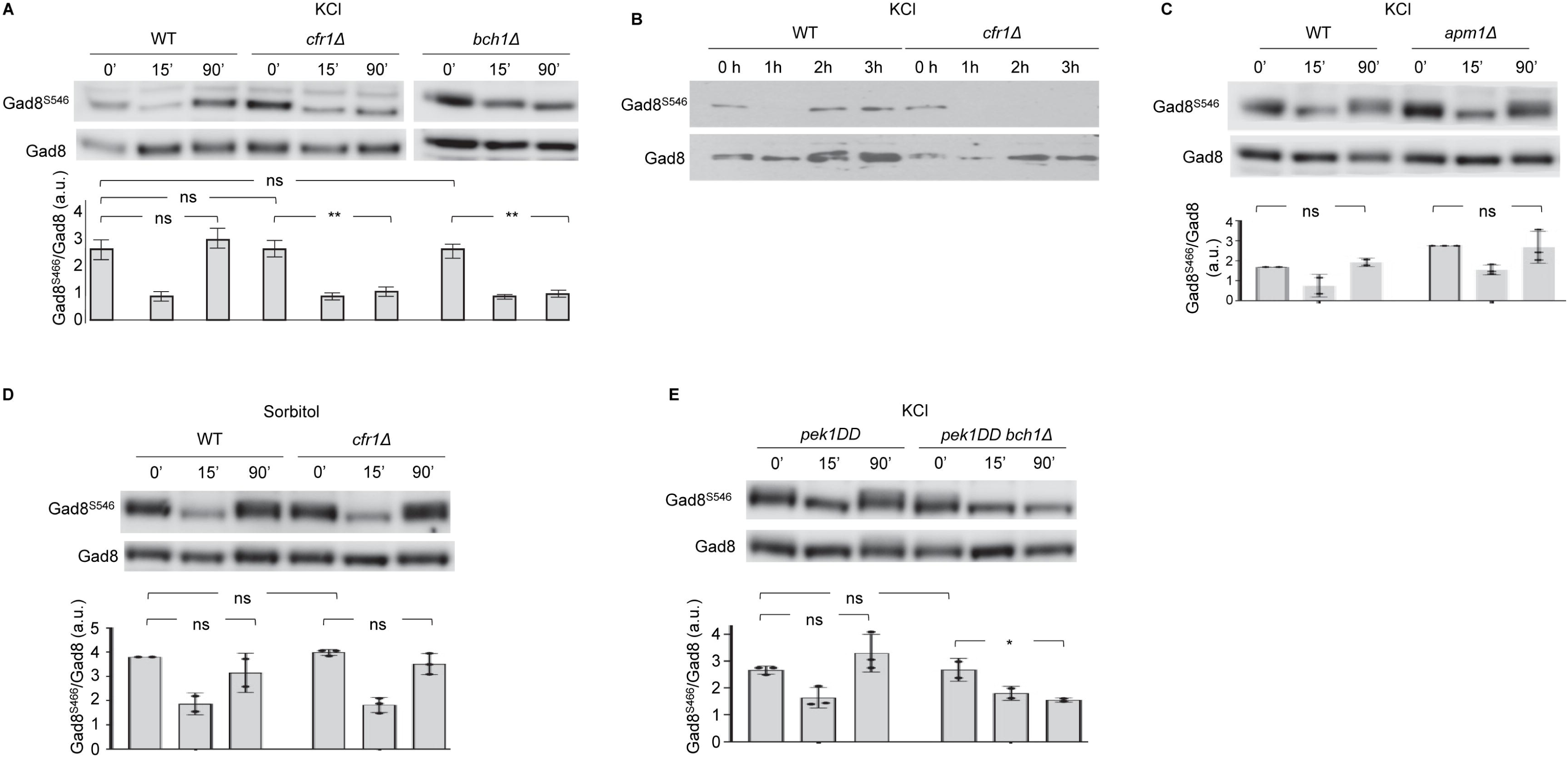
Recovery of TORC2 activity is inefficient in exomer mutants under salt stress. **(A)**. Exponentially growing cultures from the wild-type (WT) and the exomer mutants *cfr1Δ* and *bch1Δ* were treated with 1 M KCl and incubated at 28°C for the indicated times (minutes). Samples were harvested and analyzed by western blot to assess the TORC2-dependent phosphorylation of Gad8 at the Ser-546 residue (Gad8^S546^) and the total amount of Gad8 protein (Gad8). Crude cell lysates were prepared using TCA and examined by immunoblotting with specific antibodies. The graph below the blots shows the TORC2 activity calculated as the ratio of Gad8^S546^ to Gad8. The bars show the mean, standard deviation, and statistical significance determined using Tukey’s correction after ANOVA. **(B)** Same as in (A), but the samples were collected at the indicated times (hours). **(C)** Same as in (A), but the strains were the WT and the *apm1Δ* mutant, deleted for the µ subunit of the AP1 adaptor complex (Kita et al., 2004). **(D)** Same as in (A), but the cells were exposed to 1.2 M sorbitol for the indicated times (minutes). Šídák’s test was used after ANOVA. **(E)** Same as in (A), but the control and *bch1Δ* cells expressed the *pek1-S234D, T238D* (pek1DD) mutation from the *P.act1+* promoter (Moscoso-Romero et al., 2024). Šídák’s test was used after ANOVA. ns, non-significant; *, p < 0.05; **, p < 0.01. a.u., arbitrary units.

To determine whether this defect in TORC2 recovery was caused by the lack of exomer or by defective traffic from the Golgi, we evaluated the activity of TORC2 in *apm1Δ* cells, which lack the µ subunit of the AP-1 adaptor (Kita et al., 2004), treated with KCl. Based on Gad8^S546^ phosphorylation levels, *apm1Δ* was as effective as the wild-type in restoring TORC2 activity (Figure 1C).

Cells exposed to 1 M KCl undergo both osmotic stress and salt stress. Thus, we analyzed whether the lack of exomer led to a defect in the recovery of TORC2 activity in response to salt or osmotic stress. We treated the cells with 1.2 M sorbitol (osmotic stress without salt stress) and monitored the level of Gad8^S546^ phosphorylation. Figure 1D shows that the activity levels in the WT and *cfr1Δ* strains were similar over time. Thus, in the absence of exomer, cells were unable to regain TORC2 activity in response to salt, but not to osmotic stress.

We have described that exomer mutants are defective in the kinetics of CIP activity in response to osmotic stress (KCl and sorbitol), and that the defective response was independent of *tor1^+^* (Moscoso-Romero et al., 2024). To understand whether the defect in TORC2 was dependent on or independent of the CIP defect, we determined the level of Gad8^S456^ phosphorylation in *bch1^+^*and *bch1Δ* strains expressing the constitutively active Pek1 kinase under the control of the *act1^+^* promoter (*pek1DD*; Moscoso-Romero et al., 2024). As shown in Figure 1E, CIP hyperactivation by *pek1DD* in the exomer mutant did not suppress the TORC2 reactivation defect after KCl stress.

The results described above showed that the exomer mutants exhibited a defect in TORC2 reactivation after exposure to a high KCl concentration, independently of their defect in CIP activation.

### Exomer mutants exhibit defects in the Ste20^RICTOR^ localization in response to KCl

TORC2 components are located in the cell periphery (Berchtold and Walther, 2009; Tatebe et al., 2010; Ebner et al., 2017; Emmerstorfer-Augustin and Thorner, 2023), and some *S. pombe* TORC2 components and substrates bear lipid-binding and membrane-targeting domains; Sin1^Avo1^, Ste20^RICTOR^, and Gad8 bear a PH domain, an HR1 rho-binding domain, and a C2 domain, respectively. The absence of exomer led to alterations in the amount and the distribution of PM lipids (Moscoso-Romero et al., 2024). Therefore, we investigated the relationship between lipid homeostasis, TORC2 reactivation after KCl shock, and TORC2 localization

Initially, we analyzed the relationship between lipid homeostasis and Gad8^S546^ phosphorylation in response to KCl. To do so, we increased the amount of phospholipids (PLs) in exomer mutants by deleting the phosphatase gene *nem1^+^* (a regulator of the lipin phosphatidate phosphatase), which is expected to increase the level of the PL precursor phosphatidic acid, and by moderate overexpression of the Golgi-specific 1-phosphatidylinositol 4-kinase gene *pik1^+^*, the PM-specific 1-phosphatidylinositol 4-kinase gene *stt4^+^*, or the 1-phosphatidylinositol-4-phosphate 5-kinase gene *its3^+^* (Zhang et al., 2000; Park et al., 2009; Makarova et al., 2016; Willet et al., 2023; Moscoso-Romero et al., 2024). We also altered the level of sterols by expressing the C-4 methylsterol oxidase gene *erg25^+^*from the *act1^+^* promoter and by deleting the C-22 sterol desaturase gene *erg5^+^,* a deletion that reduces ergosterol synthesis and produces KCl sensitivity (Iwaki et al., 2008; Arbizzani et al., 2019; Moro et al., 2021). In all the cases, after 90 minutes of incubation in 1 M KCl, TORC2 activity was strong in the wild-type control but weak in the exomer mutants (Figure 2 A-E). These results show that increasing the amount of PLs or altering the level of sterols did not suppress the defect in restoring TORC2 activity after KCl stress.

**Figure 2.**
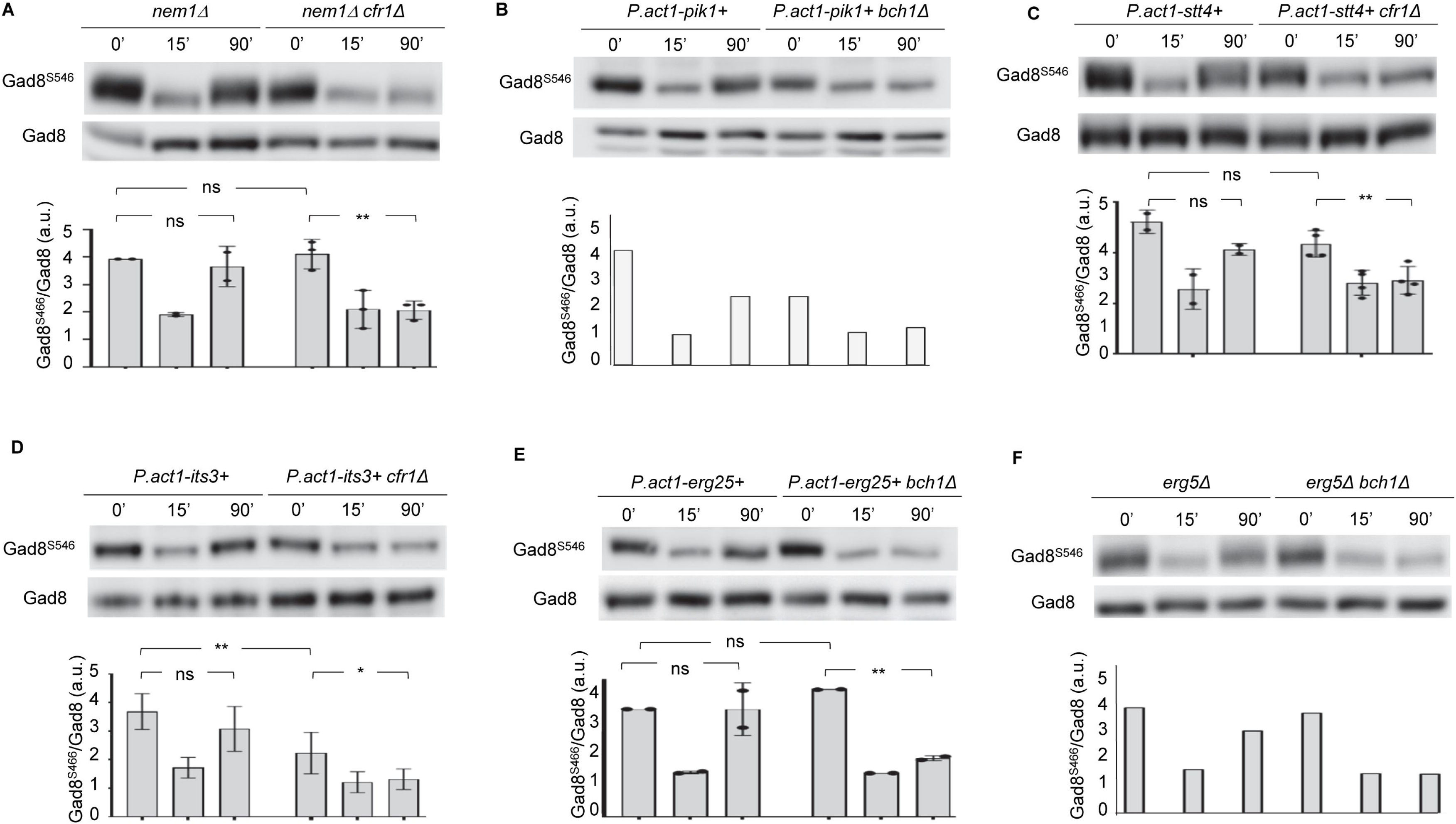
Changes in lipid homeostasis do not suppress the TORC2 activity defect of exomer mutants. **(A)** Exponentially growing cultures of the control strain and the exomer mutant *cfr1Δ* lacking the *nem1+* phosphatase were treated with 1 M KCl and incubated at 28°C for the indicated times (minutes). Samples were harvested and analyzed by western blot to assess the TORC2-dependent phosphorylation of Gad8 at the Ser-546 residue (Gad8^S546^) and the total amount of Gad8 protein (Gad8). Crude cell lysates were prepared using TCA and examined by immunoblotting with specific antibodies. The graph below the blots shows the TORC2 activity calculated as the ratio of Gad8^S546^ to Gad8. The bars show the mean, standard deviation, and statistical significance. **(B)** Same as in (A), but the exomer mutant was *bch1Δ*, and the cells overexpressed the Golgi-specific 1-phosphatidylinositol 4-kinase, type III-beta *pik1+*. **(C)** Same as in (A), but the cells overexpressed the plasma membrane 1-phosphatidylinositol 4-kinase, type III-alpha *stt4+*. **(D)** Same as in (A), but the cells overexpressed the 1-phosphatidylinositol-4-phosphate 5-kinase *its3+*. **(E)** Same as in (A), but the exomer mutant was *bch1Δ*, and the cells overexpressed the C-4 methylsterol oxidase *erg25+*. **(F)** Same as in (A), but the exomer mutant was *bch1Δ*, and the cells lacked the cytochrome P450 enzyme C-22 sterol desaturase *erg5+*. The corrections applied after ANOVA to determine the statistical significance were Šídák’s (A) and Tukey’s (C-E). ns, non-significant; *, p < 0.05; **, p < 0.01. a.u., arbitrary units. The experiments in (B) and (F) were performed only once.

Next, we analyzed the relationship between TORC2 localization and TORC2 activity using Ste20^RICTOR^-GFP as a reporter for TORC2 localization (Tatebe et al., 2010). We examined the distribution of Ste20^RICTOR^-GFP in wild-type and *cfr1Δ* cells in the absence and the presence of 1M KCl. The GFP signal was observed in both strains as a series of bright dots on the cell surface, particularly in the regions of polarized growth (cell poles and septal area), and as diffused fluorescence in the cytoplasm (Figure 3A, left panel). We observed a redistribution of the fluorescent signal 15 minutes after adding KCl, with fewer dots on the cell surface (especially on the cell poles and sides) and more dots in the cytoplasm. After 90 minutes of incubation, the Ste20^RICTOR^-GFP localization resembled that of the untreated cultures, with bright dots on the cell surface and some diffused fluorescence in the cytoplasm. Nonetheless, quantification of the number of dots in the cell surface revealed that at this time point, 30% of the wild-type cells had less than 10 dots per cell (Figure 3A, right panel), compared to 5% in the untreated culture. As for *cfr1Δ,* 85% of cells had less than 10 dots on their surface, with 25% of them exhibiting fewer than five dots. Western blot analysis showed that changes in Ste20^RICTOR^-GFP localization did not correlate with changes in the protein quantity (Figure 3B). When we incubated the cells for 15 minutes in sorbitol instead of KCl, Ste20^RICTOR^-GFP underwent a redistribution similar to that of the KCl-treated cultures. Nonetheless, after 90 minutes of incubation, more than 90% of the cells in both the wild-type and exomer mutants showed more than 10 dots on their cell surface (Figures 3A and B).

**Figure 3.**
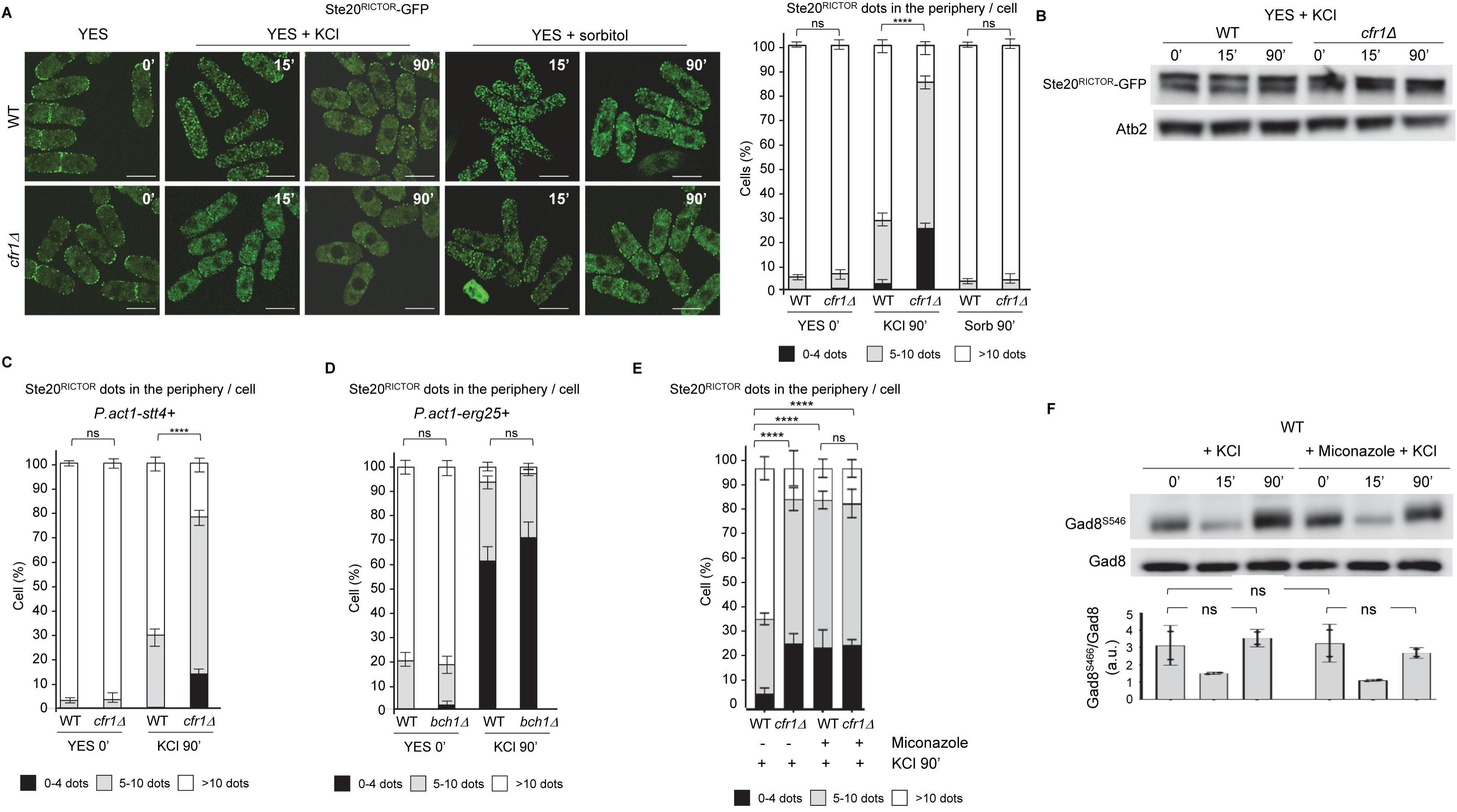
The kinetics of Ste20^RICTOR^ localization at the cell surface after KCl shock are altered in exomer mutants. **(A)** Left panel, wild-type (WT) and *cfr1Δ* cells were treated with 1 M KCl and 1.2 M sorbitol for the indicated times, and photographed under a Dragonfly confocal spinning-disc microscope. The images are average (AVG) projections. Bar, 10 µm. Right panel, the percentage of cells with Ste20^RICTOR^-GFP dots in the cell periphery was estimated from the photographs. The bars show the mean, standard deviation, and statistical significance of the differences. **(B)** Cell extracts from the wild-type (WT) and *cfr1Δ* strains bearing Ste20^RICTOR^-GFP that had been incubated in YES with 1 M KCl for the indicated times (minutes) were subjected to SDS-PAGE, and immunoblotted with anti-GFP (upper panel) and with anti-tubulin (lower panel; loading control) antibodies. **(C)** Cells from the WT and *cfr1Δ* strains that expressed Ste20^RICTOR^-GFP and overexpressed the plasma membrane 1-phosphatidylinositol 4-kinase type III-alpha *stt4+* were photographed after being treated with KCl for the indicated times. The percentage of cells with fluorescent dots in the cell periphery was estimated from the photographs. The bars show the mean, standard deviation, and statistical significance. **(D)** Same as in (C), but the exomer mutant was *bch1Δ*, and the cells overexpressed the C-4 methylsterol oxidase *erg25+*. **(E)** Same as in (C), but the cells were incubated with 1 M KCl for 90 minutes in the absence (-) and the presence (+) of miconazole. **(F)** Exponentially growing wild-type (WT) cells were treated for the indicated times (minutes) with either 1 M KCl (+KCl) or with 1 M KCl after a one-hour preincubation with miconazole (+Miconazole +KCl). Samples were harvested and analyzed by western blot to assess the TORC2-dependent phosphorylation of Gad8 at the Ser-546 residue (Gad8^S546^) and the total amount of Gad8 protein (Gad8). Crude cell lysates were prepared using TCA and examined by immunoblotting with specific antibodies. The graph below the blots shows the TORC2 activity calculated as the ratio of Gad8^S546^ to Gad8. The bars show the mean, standard deviation, and statistical significance. a.u., arbitrary units. Corrections applied after ANOVA to determine the statistical significance were Tukey’s (right panel in A for KCl, and E) and Šídák’s (right panel in A for sorbitol, C, D, and F). ns, non-significant; ****, p < 0.0001.

The results described above showed that after salt and osmotic shock, Ste20^RICTOR^ underwent a transient redistribution to the cytoplasm, and that cells lacking exomer were inefficient at relocalizing Ste20^RICTOR^ to the cell surface under salt stress. Thus, the defective reactivation of TORC2 in the exomer mutants after KCl stress appears to correlate with the compromised restoration of the Ste20^RICTOR^ localization on the cell surface.

Finally, we investigated how lipid homeostasis affected the distribution of Ste20^RICTOR^. We found that enhancing the amount of phosphatidylinositol (4,5) bisphosphate (PI-4,5-P2) by mild overexpression of the PM 1-phosphatidylinositol 4-kinase gene *stt4^+^*(Moscoso-Romero et al., 2024) did not affect the distribution and kinetics of Ste20^RICTOR^-GFP (Figure 3C). On the contrary, when we increased the amount of sterols by enhancing the expression of the *erg25^+^* gene, we found that the percentage of cells with less than 10 Ste20^RICTOR^-GFP dots on the surface was greater than that in the control cells (compare the 0’ timepoint in Figures 3A and 3D). Moreover, in cultures treated with KCl for 90 minutes, nearly all the cells had fewer than ten dots on their surface, with more than 60% having fewer than five dots, in both the wild-type and the exomer mutant. To analyze the effect of reducing sterols, we treated the cells with miconazole for 1 hour before adding 1 M KCl. After a further 90-minute incubation, the percentage of cells with less than ten Ste20^RICTOR^-GFP dots on the cell surface was above 90% in the wild-type and the mutant cells, a percentage similar to that found in the *cfr1Δ* cells treated with KCl in the absence of miconazole (Figure 3E).

These results indicated that increasing and decreasing sterol levels interfere with Ste20^RICTOR^-GFP association at the cell surface, as observed with the exomer mutants treated with KCl. However, neither mild *erg25^+^* overexpression, *erg5Δ* deletion, nor miconazole treatment altered the kinetics of Gad8^S546^ phosphorylation in the wild-type (Figures 2E, 2F, and 3F).

The results described above indicate a correlation between the presence of Ste20^RICTOR^ on the cell surface and TORC2 activity in a manner dependent on sterol homeostasis. Thus, under conditions that disrupted this homeostasis, TORC2 remained active despite most of Ste20^RICTOR^ being absent from the cell surface. Therefore, the altered kinetics of Ste20^RICTOR^ relocalization to the cell surface in the exomer mutants under salt stress may not be the only reason for their altered kinetics of Gad8^S546^ phosphorylation.

### Exomer and Ryh1^RAB6^ cooperate to activate TORC2 in response to stress

The Rab GTPase Ryh1^RAB6^ is a TORC2 activator that localizes to the Golgi and endosomes (Hengst et al., 1990; Tatebe et al., 2010; Tatebe and Shiozaki, 2010). To further study the relationship between exomer and TORC2, we investigated whether exomer functions with Ryh1^RAB6^ to regulate TORC2. First, we examined the genetic interaction between *bch1Δ* and *ryh1Δ* under different temperatures and salt stress conditions. As observed in Figure 4A, the *bch1Δ* mutation enhanced the KCl sensitivity of *tor1Δ*, *ryh1Δ*, and *ryh1-Q70L*. In addition, *bch1Δ* enhanced the thermosensitivity of the *ryh1Δ* and *ryh1-Q70L* mutants. These results suggested a functional interaction between exomer and Ryh1^RAB6^.

**Figure 4.**
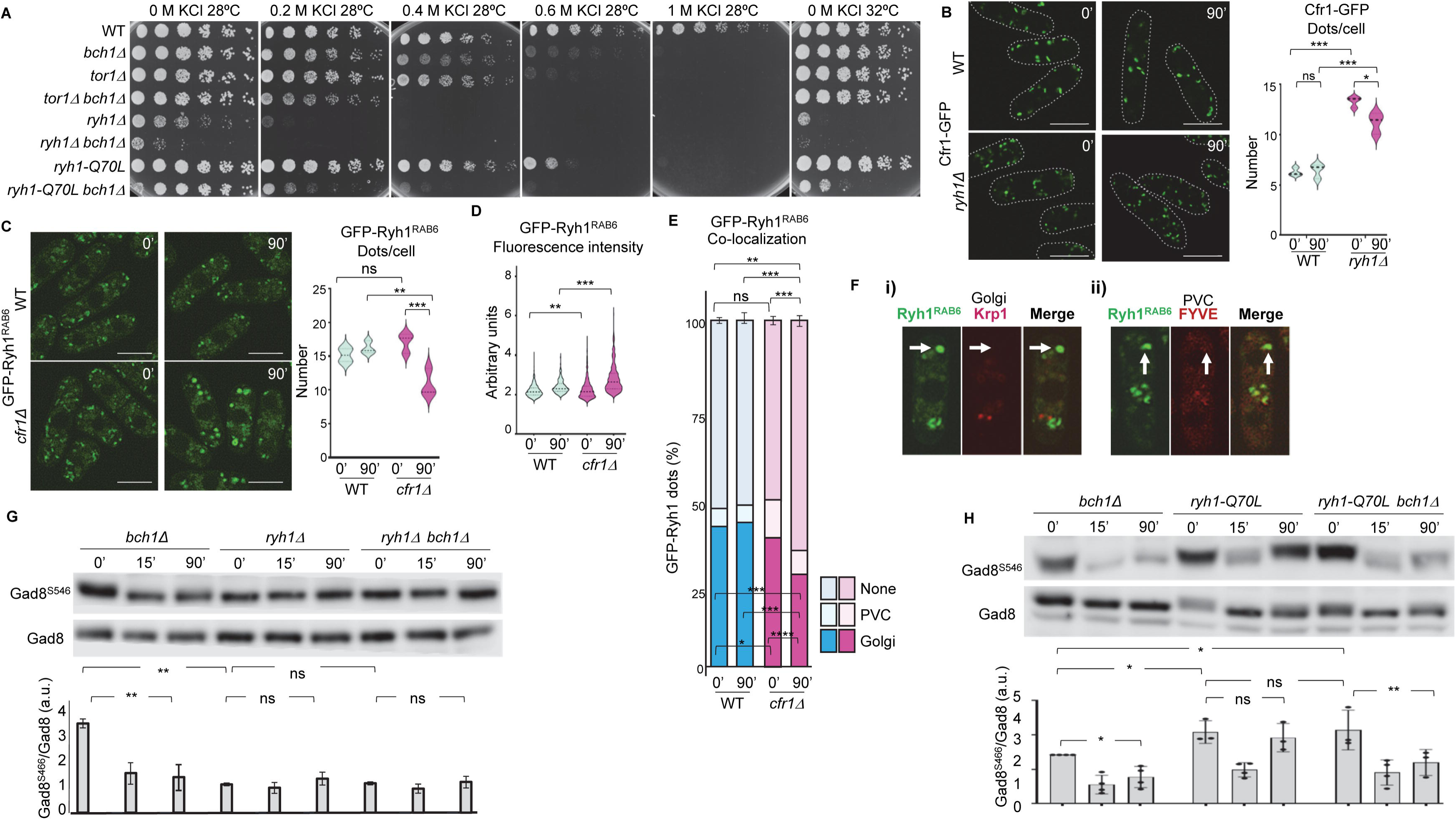
Exomer and Ryh1^RAB6^ are functionally related. **(A)** Analysis of genetic interactions. 3x10^4^ cells from the indicated strains, and serial 1:4 dilutions were inoculated on YES plates supplemented with different concentrations of KCl, and incubated at the indicated temperatures for three days. **(B)** Left panel, cells from the wild-type (WT) and *ryh1Δ* strains, bearing Cfr1-GFP, were incubated with 1 M KCl for the indicated times and photographed with a Dragonfly spinning-disc confocal microscope. Bar, 10 µm. The number of fluorescent dots was scored from the photographs and depicted in the violin graph in the right panel. **(C)** The same experimental procedure as in (B), but WT and *cfr1Δ* strains bearing GFP-Ryh1^RAB6^ were analyzed. **(D)** The fluorescence intensity of the GFP-Ryh1^RAB6^ dots scored in (C) was assessed from the photographs and depicted in a violin graph. **(E)** Co-localization analysis of GFP-Ryh1^RAB6^ with Krp1-tdTomato (Golgi marker) and Cherry-FYVE (prevacuolar compartment marker, PVC). The bar graph depicts the percentage of GFP dots that colocalized with each marker or with none of them. **(F)** Examples of cells with enlarged GFP-Ryh1^RAB6^ dots (denoted by arrows) that do not colocalize with the Golgi (i) or the PVC (ii) markers. **(G)** Exponentially growing cultures of *bch1Δ*, *ryh1Δ*, and *ryh1Δ bch1Δ*, were treated with 1 M KCl for the indicated times (minutes). Samples were harvested and analyzed by western blot to assess the TORC2-dependent phosphorylation of Gad8 at the Ser-546 residue (Gad8^S546^) and the total amount of Gad8 protein (Gad8). Crude cell lysates were prepared using TCA and examined by immunoblotting with specific antibodies. The graph below the blots shows the TORC2 activity calculated as the ratio of Gad8^S546^ to Gad8. **(H)** The same experimental procedure as in G, but the strains bore the *bch1Δ*, *ryh1-Q70L*, and *ryh1-Q70L bch1Δ* mutations. In all cases, the graphs show the mean, standard deviation, and statistical significance. Corrections applied after ANOVA to determine the statistical significance were Šídák’s (B-E) and Tukey’s (G-H). ns, non-significant; *, p < 0.05; **, p < 0.01; ***, p < 0.001; ***, p < 0.0001. a.u. arbitrary units.

Second, we analyzed whether Ryh1^RAB6^ contributed to the localization of exomer onto the Golgi/endosomes, and vice versa (He et al., 2006; Martin-Garcia et al., 2011; Hoya et al., 2017). Cfr1-GFP appeared as bright intracellular dots in both the wild-type and *ryh1Δ* strains in YES medium (0’ in the left panel of Figure 4 B) and after a 90-minute incubation in KCl. The number of Cfr1-GFP dots was greater in the *ryh1Δ* than in the *ryh1^+^* background (Figure 4B, right panel), a phenotype indicative of Golgi fragmentation, which is consistent with the role of RAB6 GTPases in maintaining the Golgi integrity (Liu and Storrie, 2015; He et al., 2006).

GFP-Ryh1^RAB6^ also appeared as bright intracellular dots in the wild-type and *cfr1Δ* cells after 0 and 90 minutes in KCl (Figure 4C, left panel). Nevertheless, the mutant cells incubated with KCl exhibited larger dots than those in the other cultures. Quantitative analyses showed that *cfr1Δ* cells treated with KCl exhibited fewer GFP-Ryh1^RAB6^ dots (right panel in Figure 4C). Additionally, the dots in the mutant were brighter than those in the control cultures (Figure 4D), which indicated an accumulation of GFP-Ryh1^RAB6^ in these structures. We performed co-localization analyses and found that in the wild-type cells incubated in YES medium, 40% of the GFP-Ryh1^RAB6^ dots co-localized with the Golgi marker Krp1-tdTomato (He et al., 2006. Figure 4E), and 10% of the dots co-localized with the prevacuolar compartment (PVC) marker mCherry-FIVE (Hoya et al., 2017). The remaining 50% of the dots did not co-localize with any marker tested, indicating the presence of Ryh1^RAB6^ in endosomes at an intermediate stage of maturation. These numbers were not significantly different when the wild-type was incubated in KCl medium for 90 minutes.

On the other hand, in the *cfr1Δ* mutant incubated in YES, there was a slight reduction in the number of GFP-Ryh1^RAB6^ dots at the Golgi, while the number of dots at the PVC increased. Furthermore, incubation of *cfr1Δ* in KCl for 90 minutes resulted in a greater reduction in the number of GFP-Ryh1^RAB6^ dots co-localized with Krp1-tdTomato, with an increase in the number of dots that did not co-localize with any marker; most of the GFP-Ryh1^RAB6^ dots that did not reside in the Golgi or the PVC were the large, bright dots (Figure 4F). In summary, the proper distribution of Ryh1^RAB6^ in the endomembranes requires exomer.

Finally, to examine if the genetic interaction between exomer and Ryh1^RAB6^ (Figure 4A) is related to TORC2 regulation, we assessed the Gad8^S546^ phosphorylation in the *bch1Δ*, *ryh1Δ*, and *bch1Δ ryh1Δ* strains. When the cells were incubated in YES (0’, Figure 4G), the phosphorylation level was lower in the *ryh1Δ* and *bch1Δ ryh1Δ* strains than in *bch1Δ*. This level was low in all the strains after 15 minutes in KCl and stayed low after 90 minutes (Figure 4G). These results showed that both with and without exomer, Ryh1^RAB6^ was required for TORC2 activity under basal and salt stress conditions. Next, we determined whether the GTP-locked form of Ryh1^RAB6^ (*ryh1-Q70L*), which renders TORC2 hyperactive (Tatebe et al., 2010), suppressed the Gad8^S546^ phosphorylation defect of *bch1Δ*. When the cells were incubated in YES, the *ryh1-Q70L* and *ryh1-Q70L bch1Δ* strains showed higher phosphorylation levels than *bch1Δ* (Figure 4H). In all strains, salt shock (15 minutes in KCl) significantly reduced the Gad8^S546^ phosphorylation level. After a 90-minute incubation in KCl, the Gad8^S546^ phosphorylation level increased in the *ryh1-Q70L* strain, but remained low in the *bch1Δ* and the *ryh1-Q70L bch1Δ* strains. Therefore, *ryh1-Q70L* did not suppress the *bch1Δ* defect in TORC2 reactivation after KCl stress.

This strongly suggests that exomer is essential for restoring the KCl-inactivated TORC2 even in the presence of the active form of Ryh1^RAB6^.

The results described above show that exomer and Ryh1^RAB6^ participate in the regulation of TORC2 activity and are both required for TORC2 reactivation in response to KCl. It is conceivable that the control of Ryh1^RAB6^ distribution by exomer is essential for TORC2 reactivation in response to salt stress.

### Efficient TORC2-Gad8 interaction requires exomer

To further investigate the relationship between exomer, Ryh1^RAB6^, and TORC2, we introduced the *bch1Δ* deletion into the hyperactive *tor1-I1816T* mutant (Halova et al., 2013). The I1816 residue is located within the FKBP12-rapamycin binding (FRB) domain of the Tor1 kinase, which is close to the catalytic domain. The FRB domain has been implicated in stress response and the interaction between the Tor kinases and some regulators (such as Avo3^RICTOR^), effectors, and substrates (Yang et al., 2013; Nguyen et al., 2018; Tafur et al., 2020). The *tor1-I1816T* mutation brings about activation of the Gad8 kinase, as indicated by the level of serine 546 phosphorylation (Halova et al., 2013). The level of Gad8^S546^ phosphorylation in *tor1-I1816T bch1^+^* and *tor1-I1816T bch1Δ* cells incubated in YES was greater than that observed in *bch1Δ* cells (0’. Figures 5A). Additionally, we observed suppression of the *bch1Δ* defect by the *tor1-I1816T* mutation in TORC2 reactivation after 90-minute KCl stress, suggesting the hyperactive Tor1 kinase can bypass the requirement of exomer in restoring the TORC2 activity after salt stress.

**Figure 5.**
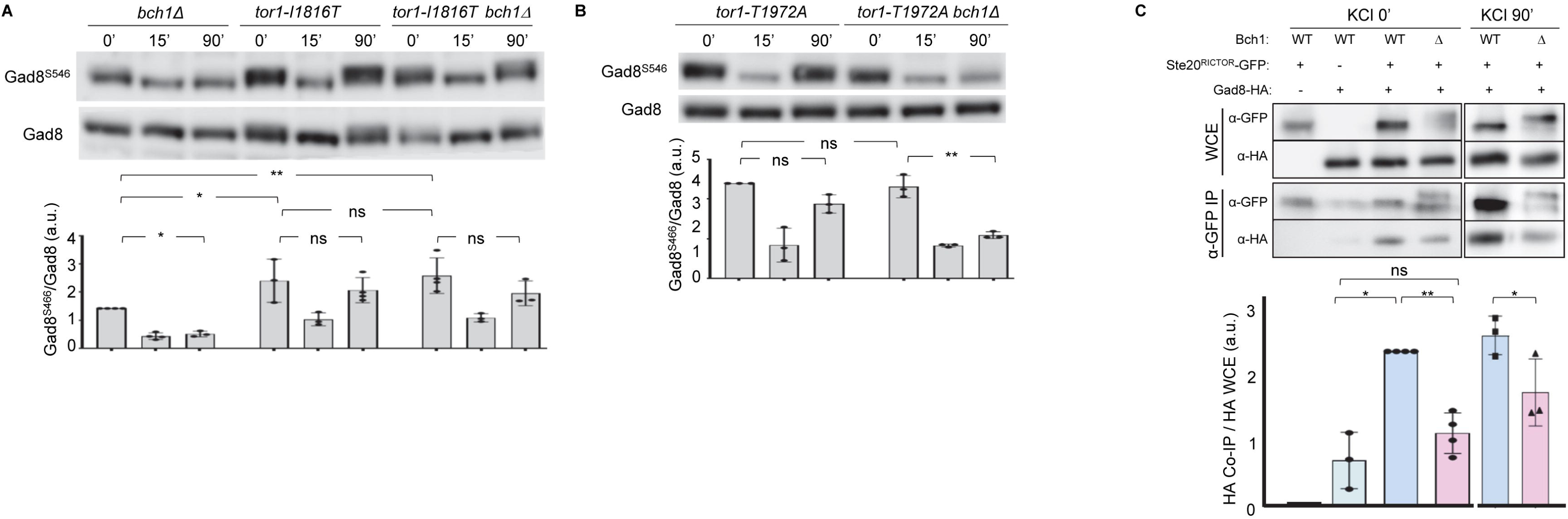
Exomer is required for efficient interaction between TORC2 and Gad8. **(A)** Exponentially growing cultures of the *bch1Δ*, *tor1-I1816T*, and *tor1-I1816T bch1Δ* mutants were treated with 1 M KCl for the indicated times (minutes). Samples were harvested and analyzed by western blot to assess the TORC2-dependent phosphorylation of Gad8 at the Ser-546 residue (Gad8^S546^) and the total amount of Gad8 protein (Gad8). Crude cell lysates were prepared using TCA and examined by immunoblotting with specific antibodies. The graph below the blots shows the TORC2 activity calculated as the ratio of Gad8^S546^ to Gad8. **(B)** The same experimental procedure as in (A), but the mutants were *tor1-T1972A* and *tor1-T1972A bch1Δ*. **(C)** Cell extracts from the wild-type (WT) and *bch1Δ* (Δ) strains, carrying Ste20^RICTOR^-GFP and/or Gad8-HA, were analyzed by western blot using anti-GFP or anti-HA monoclonal antibodies before (WCE, whole-cell extracts) or after (α-GFP IP) immunoprecipitation with anti-GFP magnetic beads. The graph below the blots depicts the ratio of the intensity of the Gad8-HA band co-precipitated with Ste20^RICTOR^ to the total HA in the cell extracts. In all the graphs, the bars show the mean, standard deviation, and statistical significance. The corrections used after ANOVA to determine the statistical significance of the differences were Tukey’s (A, C) and Šídák’s (B). ns, non-significant; *, p < 0.05; **, p < 0.01. a.u., arbitrary units.

The Tor1 T1972 residue is located within the ATP-binding domain of the kinase domain and undergoes a Gad8-dependent inhibitory phosphorylation (Halova et al., 2013). The *tor1-T1972A* mutation results in a Tor1 kinase that is insensitive to the feedback regulation by Gad8 but still susceptible to Ryh1^RAB6^ regulation. In contrast to the results obtained in *tor1-I1816T bch1Δ* cells, the level of Gad8^S546^ phosphorylation was low in *tor1-T1972A bch1Δ* cells incubated in KCl for 90 minutes, a result that showed that the *tor1-T1972A* mutation did not suppress the *bch1Δ* defect (Figure 5B). The results with both mutants indicated that, once Ryh1^RAB6^ has activated Tor1, exomer is dispensable for TORC2 reactivation.

It has been proposed that Ryh1^RAB6^ promotes the TORC2-dependent Gad8 phosphorylation by enhancing the physical interaction between TORC2 and Gad8 (Tatebe et al., 2010). We investigated whether exomer also plays a role in this interaction. To do so, we performed a co-immunoprecipitation experiment using Ste20^RICTOR^-GFP and Gad8-HA, which were expressed from their native promoters. When the cells were incubated in YES, less Gad8-HA co-precipitated with Ste20^RICTOR^-GFP in *bch1Δ* than in *bch1^+^* (Figure 5C). A similar result was obtained when the cells were incubated in KCl for 90 minutes. These results showed that exomer and Ryh1^RAB6^ share a role in facilitating or stabilizing the TORC2-Gad8 interaction.

## DISCUSSION

The Golgi apparatus plays a major role in protein and lipid trafficking between membranes and participates in signal transduction (Cancino and Luini, 2012). Exomer is a Golgi protein complex that plays a role in the response to osmotic and salt stress in budding and fission yeast (Trautwein et al., 2006; Anton et al., 2017; Hoya et al., 2017; Moro et al., 2021; Moscoso-Romero et al., 2024). *S. pombe* exomer mutants are sensitive to high KCl concentrations (Hoya et al., 2017; Moro et al., 2021). Additionally, they exhibit a defect in CIP activation in the presence of KCl because Pck2 does not associate stably with the PM due to reduced PI4P trafficking from the Golgi to the cell surface (Moscoso-Romero et al., 2024). In this work, we found that TORC2 reactivation after exposure to high KCl concentrations is compromised in the exomer mutants. There is cross-regulation between CIP and TORC2 (Madrid et al., 2015; Du et al., 2016; Madrid et al., 2016; Cansado et al., 2021). Thus, CIP activity reduces TORC2 signaling in response to certain stressors. Based on this information, the reduced CIP activity of exomer mutants should result in increased TORC2 activity, which contradicts our findings. Additionally, the inability of constitutively activated CIP to suppress the TORC2 defect, together with the fact that CIP is deficient in response to KCl and sorbitol, while TORC2 efficiently responds to sorbitol suggest that the CIP and TORC2 signaling defects of the exomer mutants are distinct and independent.

### Ste20^RICTOR^ localization in response to stress

Lipid homeostasis and distribution are crucial for cell physiology (Santos and Preta, 2018). Several PM microdomains, enriched in specific lipids and proteins, have been described in fungi. TORC2 accumulates into the MCT (Membrane compartments of TOR) microdomains, which are adjacent to but distinct from MCC (membrane compartments of Can1), also known as eisosomas (Berchtold and Walther, 2009; Athanasopoulos et al., 2019). Eisosomes are microdomains enriched in sterols and some proteins that bind phosphatidylinositol 4, 5-bisphosphate (PI(4,5)P2). Eisosomes, sterols, and PI(4,5)P2 play significant roles in PM homeostasis, stress response, and TORC2 signaling (Berchtold and Walther, 2009; Kabeche et al., 2015; Madrid et al., 2016; Riggi et al., 2018; Roelants et al., 2018; Cansado et al., 2021; Sakata et al., 2022; Thorner, 2022). Several TORC2 components have lipid-binding domains that promote the anchoring of the complex to the PM. However, some of these domains are dispensable for the TORC2 activity, and some lipids play a regulatory, rather than a structural, role in TORC2 regulation (Tatebe et al., 2010; Tatebe et al., 2017; Thorner, 2022; Emmerstorfer-Augustin and Thorner, 2023). In *S. cerevisiae*, changes in membrane tension produced by osmolality fluctuations modulate TORC2 activity by altering the distribution of some of its components and substrates in the PM microdomains (Berchtold and Walther, 2009; Riggi et al., 2018; Roelants et al., 2018). Although *S. pombe* exomer mutants exhibit defects in the distribution of sterols and PI(4,5)P2 (Moro et al., 2021; Moscoso-Romero et al., 2024), changes in the levels of these lipids did not complement the TORC2 defect. This result suggests that the distribution rather than the levels of these lipids may modulate TORC2. According to Ste20^RICTOR^-GFP localization, osmotic shock leads to the internalization of *S. pombe* TORC2 and its distribution in internal structures whose appearance differs from that of the Golgi and endosomes. An intriguing possibility is that they are stress granules (SGs). These structures modulate the response of several signaling pathways to stress by sequestering or bringing together some of their components (Takahara and Maeda, 2012; Cadena Sandoval et al., 2021; Lee et al., 2025). Further studies will be required to address this point and the involvement of exomer in the process.

After adaptation, Ste20^RICTOR^-GFP relocalizes to the cell surface, a process that resembles the recovery of Gad8^S546^ phosphorylation in both the wild-type and exomer mutants. These results suggest that the presence of TORC2 on the PM is required for its activity. Nevertheless, the fact that variations in sterol homeostasis reduced Ste20^RICTOR^-GFP localization on the cell periphery, even in the presence of exomer and the absence of stress, argued against Ste20^RICTOR^ localization on the PM being required for TORC2 activity. Although it is possible that the observed changes in the distribution of Ste20^RICTOR^-GFP did not reflect changes in the distribution of the entire complex, these results suggest that TORC2 may be activated in other locations. We could not determine whether the substrate followed the same kinetics and where the complex interacted with the substrate, because fusing Gad8 to GFP in its N- and C-terminal ends did not produce a specific fluorescent signal on the cell surface or internal structures either in the absence or presence of stress (Tatebe et al., 2010 and our unpublished results).

### Relationship between exomer, Ryh1^RAB6^, and TORC2 regulation in response to KCl

Ryh1^RAB6^ and Ypt3^RAB11^ are GTPases that localize in the Golgi/endosomes and, as exomer, participate in secretion, and are required for efficient TORC2 response (Hengst et al., 1990; He et al., 2006; Ma et al., 2010; Tatebe et al., 2010; Tatebe and Shiozaki, 2010; Cheng, 2002 #6668; Hoya et al., 2017; Moro et al., 2021; Moscoso-Romero et al., 2024). Therefore, exomer may be functionally related to Ryh1^RAB6^ and/or Ypt3^RAB11^ in terms of TORC2 regulation. The observed genetic interaction between *bch1+* and *ryh1^RAB6+^* supports this hypothesis. The fact that *apm1Δ* exhibits a TORC2 response similar to that of the wild type a priori rules out the hypothesis that the deficient response in the *ryh1Δ*, *ypt3-i5*, and exomer mutants is an indirect consequence of impaired vesicle trafficking.

The absence of exomer leads to partial Ryh1^RAB6^ mislocalization, suggesting that this defect may be the leading cause or a contributing factor for reduced Gad8^S546^ phosphorylation. *ryh1-Q70L* cannot suppress the defect in Gad8^S546^ phosphorylation in *bch1Δ*, a result that suggests that this mislocalization is relevant to the low TORC2 activity in exomer mutants. TORC2 is mainly localized at the cell surface, where a fraction of the GTP-locked GTPase has been observed, a result that suggests that this fraction activates TORC2 in the PM (He et al., 2006; Ma et al., 2010; Tatebe et al., 2010). Nevertheless, determining the localization of Ryh1-Q70L in exomer mutants will help to understand the relationship between exomer, Ryh1^RAB6^ localization, and TORC2 activation in response to KCl. Ryh1^RAB6^ activates TORC2, but it is not essential for the assembly of the complex. This led to the conclusion that the GTPase facilitates the interaction between the complex and its substrate (Tatebe et al., 2010). In exomer mutants, the TORC2-Gad8 binding is reduced. This finding is consistent with the idea that the exomer plays a role in regulating TORC2 through a mechanism related to Ryh1^RAB6^.

*ryh1^RAB6+^* interacts genetically with *ypt3^RAB11+^*, and TORC2 activity is low in the *ypt3-i5* strain. Nevertheless, GTP-locked Ypt3Q69L does not activate TORC2 (He et al., 2006), indicating that this GTPase is not a TORC2 activator. In the *ypt3-i5* mutant, as in exomer mutants, there are defects in the morphology of the Golgi and endosomes (Hoya et al., 2017; Moro et al., 2021, Cheng, 2002 #6668) and Ryh1^RAB6^ is partially delocalized such that part of this protein does not co-localize with endosomal markers (FM4-64 9 and mCherry-FIVE in the *ypt3-i5* and the exomer mutants, respectively). These results support the hypothesis that Ryh1 localization is important for TORC2 activation (He et al., 2006). *ryh1-Q70L* suppresses the TORC2 activity defect of *ypt3-i5* but not that of *bch1Δ*. This could be because Ypt3^RAB11^ and exomer act through different mechanisms, or because *ypt3-i5* is a point mutation and *bch1Δ* is a deletion. Further studies are necessary to determine whether there is a functional relationship between exomer and Ypt3^RAB11^.

As discussed above, the presence of Ste20^RICTOR^ in the PM correlates with TORC2 activation, unless the level of sterols is altered. This supports the notion that either these lipids or an adequate lipid environment in the PM contribute to TORC2 regulation. Lipid environment affects the distribution and activity of proteins involved in signaling, both in the PM and internal organelles (Mayinger, 2009; 2011; Stone et al., 2017; Sunshine and Iruela-Arispe, 2017; Baccouch et al., 2022; Sarmento et al., 2023). Golgi cisternae and endosomes undergo maturation processes, including changes in the lipid composition of their membranes that are important for their functionality (Losev et al., 2006; Huotari and Helenius, 2011; Scott, 2011 #5933; Daboussi et al., 2012; Scott et al., 2014). Exomer mutants have defects in the level and trafficking of PI4P, and in the integrity of the Golgi cisternae and endosomes (Hoya et al., 2017; Moro et al., 2021; Moscoso-Romero et al., 2024). We propose that these defects cause Ryh1^RAB6^ to be located in an environment that prevents it from facilitating the contact between TORC2 and Gad8. This would not be a general mechanism of TORC2 regulation, because under glucose starvation, Gad8^S546^ phosphorylation recovers over time independently of Ryh1^RAB6^ (Hatano et al., 2015). TORC2 regulation in response to glucose appears to be related to the PI(4,5)P2 synthesis in the PM (Li et al., 2014; Madrid et al., 2016). The Ryh1^RAB6^-dependent regulation in response to KCl may be related to changes in PM tension and/or microdomain organization, which would then be transduced to the endomembrane system. In the absence of exomer, these changes would be more drastic, altering the Ryh1^RAB6^ distribution and, consequently, the regulation of TORC2.

The defect in TORC2 reactivation, together with the defects in CIP signaling in response to osmotic stress and the distribution of the K^+^ and Ca^2+^ channels and pumps, would contribute to the potassium sensitivity of exomer mutants (Hoya et al., 2017; Moro et al., 2021; Moscoso-Romero et al., 2024).

## ACKNOWLEDGEMENTS

We thank J. Cansado, T. Kuno, J. Petersen, S. Moreno, P. Pérez, Y. Sanchez, and the YGRC (http://yeast.lab.nig.ac.jp) for strains and plasmids. Financial support from the University of Salamanca (grant 18 K261-463AC01) and from the Spanish Ministerio de Ciencia e Innovación (MCIN/AEI/10.13039/501100011033/ FEDER “Una manera de hacer Europa”) to MH Valdivieso (grant PID2020-115111GB-I00), and from the Junta de Castilla y Leon/European Union FEDER program (grants “Escalera de Excelencia” CLU-2017-03/14-20 and “Internationalization Project CL-EI-2021-08) to the IBFG made this work possible. EMR and SM were supported by predoctoral fellowships from the Junta de Castilla y León and the Spanish Ministerio de Educación, respectively.

---

**Table S1.** List of yeast strains used in this work.

| STRAIN | GENOTYPE | SOURCE |
| --- | --- | --- |
| HVP30 | *leu1-32 his3-Δ1 ura4-Δ18 ade6 h-* | Lab stock |
| HVP264 | *cfr1Δ::his3+ leu1-32 his3-Δ1 ura4-Δ18 ade6 h-* | Lab stock |
| HVP1733 | *pmk1+::pmk1-HA6His:ura4+ ura4-Δ18 leu1-32 ade6 h+* | J Cansado |
| HVP2357 | *bch1Δ::KANMX6 leu1-32 his3-Δ1 ura4-Δ18 ade6 h-* | Lab stock |
| HVP3490 | *apm1Δ::ura4 leu1-32 his3-Δ1 ura4-Δ18 ade6 ?h-* | T. Kuno |
| HVP3712 | *cfr1Δ::his3+ leu1+::cfr1-GFP leu1-32 his3-Δ1 ura4-Δ18 ade6 h-* | Lab stock |
| HVP4068 | *HPHMX6 (hph.171k) leu1-32 his3-Δ1 ura4-Δ18 h+* | Lab stock |
| HVP4724 | *gad8::ura4+ gad8+::gad8-6HA:KANMX6 leu1-32 ura4-D18 ade6-M210 h90* | YGRC |
| HVP4734 | *tor1::KANMX6 h+* | S. Moreno |
| HVP4757 | *ryh1::KANMX6 leu1-32 h-* | CA6217, Shiozaki lab stock |
| HVP4758 | *ryh1::FLAG-ryh1-Q70L leu1-32 h-* | CA6817, Shiozaki lab stock |
| HVP4763 | *tor1::KANMX6 bch1::NATMX6 h+* | This work |
| HVP4786 | *ryh1::FLAG-ryh1-Q70L bch1::NATMX6 leu1-32 h-* | This work |
| HVP4796 | *ryh1::KANMX6 bch1::NATMX6 leu1-32 h-* | This work |
| HVP4813 | *ryh1::KANMX6 leu1+::GFP-ryh1 leu1-32 h-* | This work |
| HVP4893 | *ryh1::KANMX6 leu1+::GFP-ryh1 cfr1::his3+ leu1-32 his3-Δ1? ura4-Δ18? ade6? h?* | This work |
| HVP4992 | *ste20+::ste20-3xGFP:KANMX6 leu1-32 h-* | CA6655 Shiozaki lab stock |
| HVP5021 | *ste20+::ste20-3xGFP:KANMX6 bch1::NATMX6 leu1-32 h-* | This work |
| HVP5026 | *erg5::KANMX6 leu1-32 ura4-Δ18 ade6 h-* | P. Perez |
| HVP5446 | *erg5::KANMX6 bch1::NATMX6 leu1-32 ura4-Δ18 ade6 h-* | This work |
| HVP5600 | *hph171k::HPHMX6:P.act1-pek1-S234D,T238D:NATMX6 pmk1+::pmk1-HA6His:ura4+ leu1-32 his3-Δ1 ura4-Δ18 ade6? h+* | Lab stock |
| HVP5601 | *hph171k::HPHMX6:P.act1-pek1-S234D,T238D:NATMX6 bch1::NATMX6 pmk1+::pmk1-HA6His:ura4+ leu1-32 his3-Δ1 ura4-Δ18 ade6? h+* | Lab stock |
| HVP5692 | *tor1-T1972A h-* | JP1560. J Petersen |
| HVP5693 | *tor1.I1816T h+* | JP1563 J Petersen |
| HVP5719 | *tor1-T1972A pmk1+::pmk1-HA6His:ura4+ h?* | Lab stock |
| HVP5735 | *its3+::HPHMX6:P.act1-its3+ leu1-32 his3-Δ1 ura4-Δ18 ade6 h-* | Lab stock |
| HVP5738 | *tor1-I1816T pmk1+::pmk1-HA6His:ura4+ h?* | Lab stock |
| HVP5747 | *its3+::HPHMX6:P.act1-its3+ bch1::KANMX6 leu1-32 his3-Δ1 ura4-Δ18 ade6 h-* | Lab stock |
| HVP5762 | *tor1-T1972A pmk1+::pmk1-HA6His:ura4+ bch1::KANMX6 h?* | Lab stock |
| HVP5763 | *tor1-I1816T pmk1+::pmk1-HA6His:ura4+ bch1::KAN h?* | Lab stock |
| HVP5868 | *stt4+::HPHMX6:P.act1-stt4+ leu1-32 his3-Δ1 ura4-Δ18 ade6 h-* | Lab stock |
| HVP5887 | *stt4+::HPHMX6:P.act1-stt4+ cfr1::his3+ leu1-32 his3-Δ1 ura4-Δ18 ade6 h-* | Lab stock |
| HVP5927 | *nem1::NATMX6 leu1-32 his3-Δ1 ura4-Δ18 ade6 h+* | Lab stock |
| HVP5728 | *nem1::NATMX6 cfr1::his3+ leu1-32 his3-Δ1 ura4-Δ18 ade6 h-* | Lab stock |
| HVP5944 | *stt4+::HPHMX6:P.act1-stt4+ ste20+::ste20-3xGFP:KANMX6 leu1-32 his3-Δ1? ura4-Δ18? ade6? h-* | This work |
| HVP5957 | *stt4+::HPHMX6:P.act1-stt4+ ste20+::ste20-3xGFP:KANMX6 bch1::NATMX6 leu1-32 his3-Δ1? ura4-Δ18? ade6? h-* | This work |
| HVP6017 | *gad8::ura4+ gad8+::gad8-6HA:KANMX6 ste20+::ste20-3xGFP:KANMX6 leu1-32 ura4-D18? ade6? h-* | This work |
| HVP6018 | *gad8::ura4+ gad8+::gad8-6HA:KANMX6 ste20+::ste20-3xGFP:KANMX6 bch1::NATMX6 leu1-32 ura4-D18? ade6? h-* | This work |
| HVP6090 | *erg25+::HPHMX6:P.act1-erg25+ leu1-32 his3-Δ1 ura4-Δ18 ade6 h+* | This work |
| HVP6093 | *erg25+::HPHMX6:P.act1-erg25+ bch1::NATMX6 leu1-32 his3-Δ1 ura4-Δ18 ade6 h+* | This work |
| HVP6094 | *erg25+::HPHMX6:P.act1-erg25+ ste20+::ste20-3xGFP:KANMX6 leu1-32 his3-Δ1? ura4-Δ18? ade6? h?* | This work |
| HVP6095 | *erg25+::HPHMX6:P.act1-erg25+ ste20+::ste20-3xGFP:KANMX6 bch1::NATMX6 leu1-32 his3-Δ1? ura4-Δ18? ade6? h?* | This work |
| HVP6109 | *HPHMX6 (hph.171k):P.act1-pik1+ pmk1+::pmk1-HA6His:ura4+ leu1-32 his3-Δ1 ura4-Δ18 h+* |  |
| HVP6115 | *HPHMX6 (hph.171k):P.act1-pik1+ pmk1+::pmk1-HA6His:ura4+ bch1::KANMX6 leu1-32 his3-Δ1 ura4-Δ18 h+* |  |

## Notes

### Competing Interest Statement

The authors have declared no competing interest.

